# The gut microbiota is essential for *Trichinella spiralis*- evoked suppression of colitis

**DOI:** 10.1101/2024.04.09.588742

**Authors:** Hualei Sun, Shao Rong Long, Miao Jiang, Hui Ran Zhang, Jing Jing Wang, Zi Xuan Liao, Jing Cui, Zhong Quan Wang

## Abstract

Inflammatory bowel disease (IBD) increases the risk of colorectal cancer, and it has the potential to diminish the quality of life. Clinical and experimental evidence demonstrate protective aspects of parasitic helminth infection against IBD. However, studies on the inhibition of inflammation by helminth infection have overlooked a key determinant of health: the gut microbiota. Infection with helminths induces alterations in the host microbiota composition. However, the potential influence and mechanism of helminth infections induced changes in the gut microbiota on the development of IBD has not yet been elucidated. In this study, we analyzed the intersection of helminth *Trichinella spiralis* and gut bacteria in the regulation of colitis and related mechanisms. *T. spiralis* infected mice were treated with antibiotics or cohused with wild type mice, then challenged with DSS-colitis and disease severity, immune responses and goblet cells assessed. Gut bacteria composition was assessed by 16 s rRNA sequencing and SCFAs were measured. Results showed that protection against disease by infection with *T. spiralis* was abrogated by antibiotic treatment, and cohousing with *T. spiralis*- infected mice suppressed DSS-colitis in wild type mice. Bacterial community profiling revealed an increase in the abundance of the bacterial genus *Muribaculum* and *unclassified_Muribaculaceae* in mice with *T. spiralis* infection or mice cohoused with *T. spiralis*- infected mice. Metabolomic analysis demonstrated increased propionic acid in feces from *T. spiralis*- infected mice. Data also showed that the gut microbiome modulated by *T. spiralis* exhibited enhanced goblet cell differentiation and elevated IL-10 levels in mice. Taken together, these findings identify the gut microbiome as a critical component of the anti-colitic effect of *T. spiralis* and gives beneficial insights into the processes by which helminth alleviates colitis.

**Author Summary:** Inflammatory bowel disease (IBD) encompasses Crohn’s Disease and Ulcerative Colitis. It affects both children and adults. Reports have highlighted the potential use of helminths or their byproducts as a possible treatment for IBD. Accumulating evidence also suggests that the gut microbiota is a key factor in modulating IBD. In this study, we revealed the protective effect of a prior infection with *T. spiralis* on DSS-induced colitis in mice. Specifically, *T. spiralis* infection reshaped the gut microbiome of mice, resulting in an increased abundance of SCFA-producing bacteria *Muribaculum* and *unclassified_Muribaculaceae* and thereby producing a larger amount of propionic acid. Furthermore, the gut microbiome modulated by *T. spiralis* exhibited enhanced goblet cell differentiation and elevated IL-10 levels, ultimately ameliorating experimental colitis. These findings suggest that the modulation of host microbiota during *T. spiralis* infection plays a crucial role in the suppression of colitis, and any intention-to-treat with helminth therapy should be based on the patient’s immunological and microbiological response to the helminth.

## Introduction

Inflammatory bowel disease (IBD) encompasses Crohn’s Disease and Ulcerative Colitis. It affects both children and adults. The symptoms of the IBD are mild to severe and may threaten life, which not only include fever and diarrhea, but also abscess formation, stenosis, and the development of colitis-associated colorectal cancer [1, 2]. Multiple etiological factors, such as environmental factors, genetic background, and dysregulation of the immune system, are involved in IBD [3]. Since the middle of the twentieth century, the incidence of IBD has increased in the Western world, and now it has emerged in newly industrialized countries in Asia, South America and Middle East and has evolved into a global disease with rising prevalence in every continent [4].

Epidemiological studies have reported that within the last century helminths have gone from being ubiquitous to all but absent in developed countries. Further epidemiological investigations demonstrated that IBD is less prevalent in helminth-endemic countries [5]. Several different types of helminth infection in mice colitis models has also been shown some species, such as *Schistosoma mansoni* [6]*, Heligmosomoides polygyrus* [7]*, Trichinella spiralis* [8] and *Hymenolepis diminuta* [9], ameliorated the inflammatory reaction demonstrating certain anti-colitic effects. Most of these responses were characterized by an increase in Th2 immune response and regulatory T cells (Tregs), which consequently result in the secretion of regulatory cytokines that have anti-inflammatory characteristics.

In addition to being associated with immune cells, the gut microbiota is considered a crucial environmental factor in IBD [10]. Accumulating evidence suggests that the gut microbiota is a key factor in modulating the host immune system, influencing a predisposition to autoimmune diseases, including IBD [11]. A range of bacterial species, including *Lactobacillus, Bifidobacterium*, and *Faecali bacterium*, have shown this protective role via up-regulation of IL-10 production and down-regulation of pro-inflammatory cytokines. Moreover, *Clostridium and Bacteroides* species induced the expansion of Tregs to mitigate intestinal inflammation [12].

Helminths coexist with the gut microbiota and their mammalian hosts, and can induce changes in the composition of the host’s gut microbiome [13]. However, the potential influence and mechanism of helminth infections inducing alterations in the gut microbiota on the development of IBD has not yet been elucidated. In this study, we analyzed the intersection of helminth *T. spiralis* and gut bacteria in the regulation of colitis and related mechanisms. The data herein, show that the modulation of host microbiota by *T. spiralis* is essential to the suppression of colitis.

## Results

### Antibiotics treatment abrogates *T. spiralis*-evoked suppression of colitis

The possibility that the gut bacteria participated in *T. spiralis*-evoked suppression of colitis was tested with broad-spectrum antibiotics (**Fig 1A**). Data showed that mice present antibiotics (ABX) treatment during the induction of colitis exhibited an earlier onset of rectal bleeding and diarrhea compared to mice without ABX treatment (**S1 Fig**). As expected, the DSS-treated group exhibited visible signs of inflammation characterized by body weight loss, rectal bleeding, and diarrhea, leading to a significantly increased level of disease activity index (DAI) (**Fig 1B and 1C**), typical pathological changes, including epithelial erosion, edema, loss of the mucus layer, substantial polymorphonuclear infiltrate into the lamina propria, and increased histopathological score, and pre-infection of *T. spiralis* led to a significant decrease in weight loss and disease symptoms in DSS colitis (**Fig 1D and 1E**). However, the suppression of DSS-induced colitis evoked by infection with *T. spiralis* was absent in mice co-treated with ABX (**Fig 1B-1E**). The control, *T. spiralis* infection and ABX treatment alone mice showed no weight loss or microscopic damage of the colon. These results demonstrate that the gut bacteria participated in *T. spiralis*-evoked suppression of colitis.

**Fig 1.**
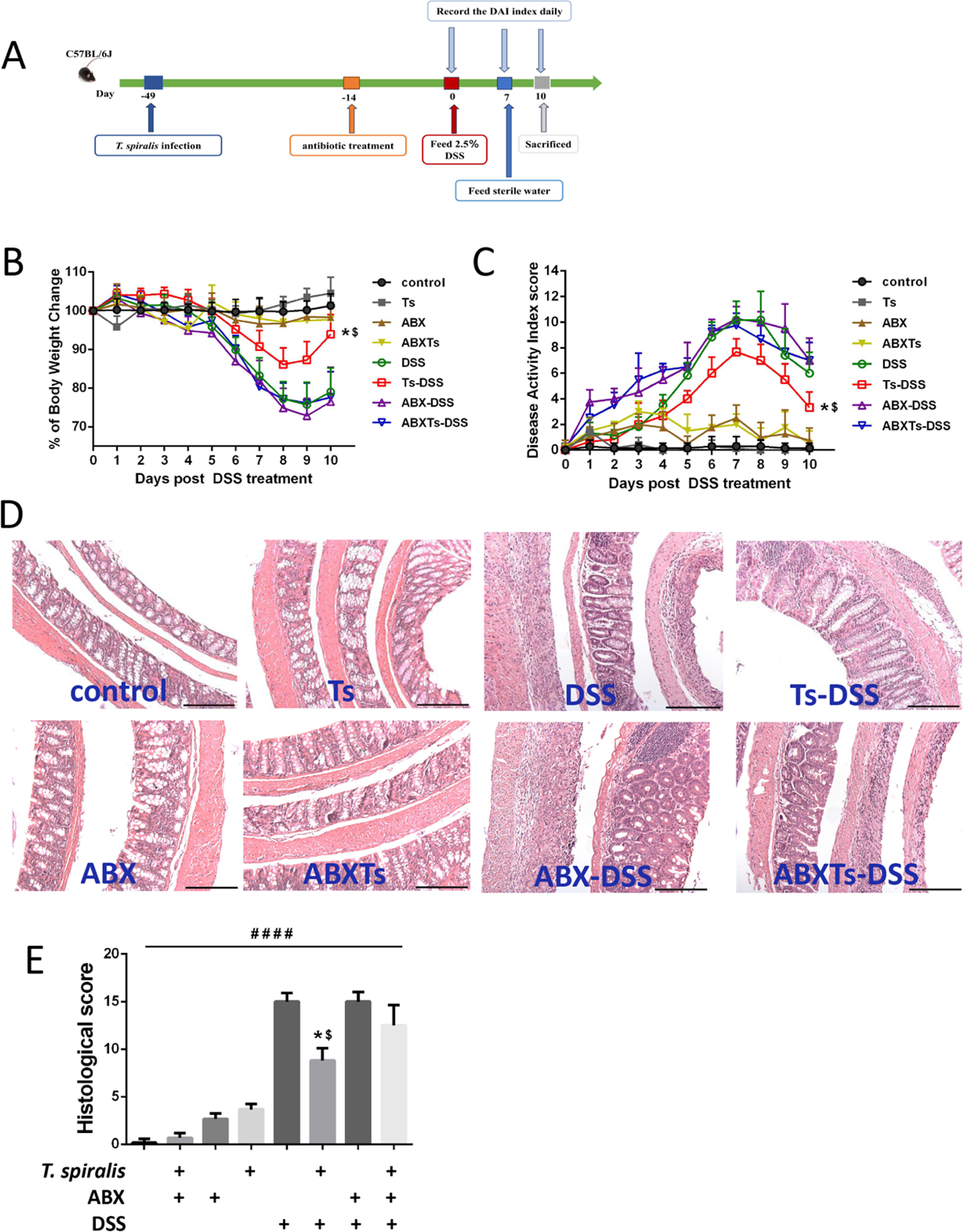
Broad-spectrum antibiotic treatment prevents *T. spiralis*-evoked inhibition of colitis. (**A**) *T. spiralis* infection, ABX treatment and DSS-induced colitis schedule, mice were orally gavaged with *T. spiralis* or PBS, 35 days later, some of the mice were treated with ABX daily for 14 days, then were given drinking water containing 2.5% (wt/vol) DSS ad libitum for 7 days, and others left untreated as controls. After a further 3 days of distilled water fed, mice were sacrificed and samples were obtained and processed. (**B**) Weight change in percent. (**C**) The changes in disease activity index (DAI), scored from diarrhea, bleeding and body weight loss. (**D**) The histopathological changes in the colon tissues were examined by H&E staining, the black bar indicates 200 μm. (**E**) Histopathological scores were determined for the colon tissue samples. The data shown are means ± SD. Representative results from one out of three independent experiments with n = 5. *, $ *P*< 0.05 compared to DSS and ABXTs-DSS, respectively; ####*P* < 0.0001 versus the respective control group. ABX: antibiotic cocktail of ampicillin (1g/L), vancomycin (0.5g/L), neomycin (1g/L), and metronidazole (1g/L) treated; ABXTs: *T. spiralis* infected and antibiotic treated; Ts: *T. spiralis* infected; DSS: DSS-induced colitis; Ts-DSS: *T. spiralis* infected and DSS induced colitis; ABX-DSS: antibiotic treated and DSS induced colitis; ABXTs-DSS: *T. spiralis* infected, antibiotic treated and DSS induced colitis

### *T. spiralis*- infected mice alters gut microbiome composition

The influence of *T. spiralis* infection on mouse gut microbiota was determined using bacterial 16S rRNA gene sequencing. Venn diagram displayed decrease in operational taxonomic units (OTUs) in mice-infected with *T. spiralis* (**Fig 2A**). The Chao1 index and ACE index of α diversity detection showed that *T. spiralis*- infected mice had a lower diversity of microbiota than control mice (**Fig 2B**). Besides, β diversity analysis of Principal Co-ordinates Analysis (PCoA) based on Binary-Jaccard demonstrated that distinct clustering of microbiota composition between *T. spiralis*- infected mice and the control mice (**Fig 2C**). These data suggested that *T. spiralis* infection changed gut microbiota diversity and composition in mice. Then, we analyzed the bacterial profiles at the phylum and genus levels to further illuminate the difference of microbiota composition. At phylum level, Bacteroidota and Firmicutes were the dominant phyla followed by Proteobacteria, Patescibacteria, Actinobacteriota, Desulfobacteriota, Campylobacterota, Verrucomicrobiota and Defferribacterota (**Fig 2D**). And data showed that *T. spiralis* infection increased the relative abundance of Bacteroidota and decreased the relative abundance of Firmicutes (**Fig 2D**). In terms of bacterial composition, our results showed that *T. spiralis* infection induced significant changes in gut microbiome composition, characterized by an increase in the abundance of the bacterial genus *Muribaculum* and *unclassified_Muribaculaceae* and a decrease in the abundance of the bacterial genus *unclassified_Oscillospiraceae* and *Alistipes* in the top 20 genera (**Fig 2E and 2F**). In addition to taxonomic composition, the functional profiles of microbial communities were predicted using the PICRUSt2 software based on 16S rRNA gene-based microbial compositions. Significant differences were detected in KEGG pathways between the two groups, *T. spiralis* infection enhanced the function of lipid metabolism (*P*=0.0416) (**Fig 2G**).

**Fig 2.**
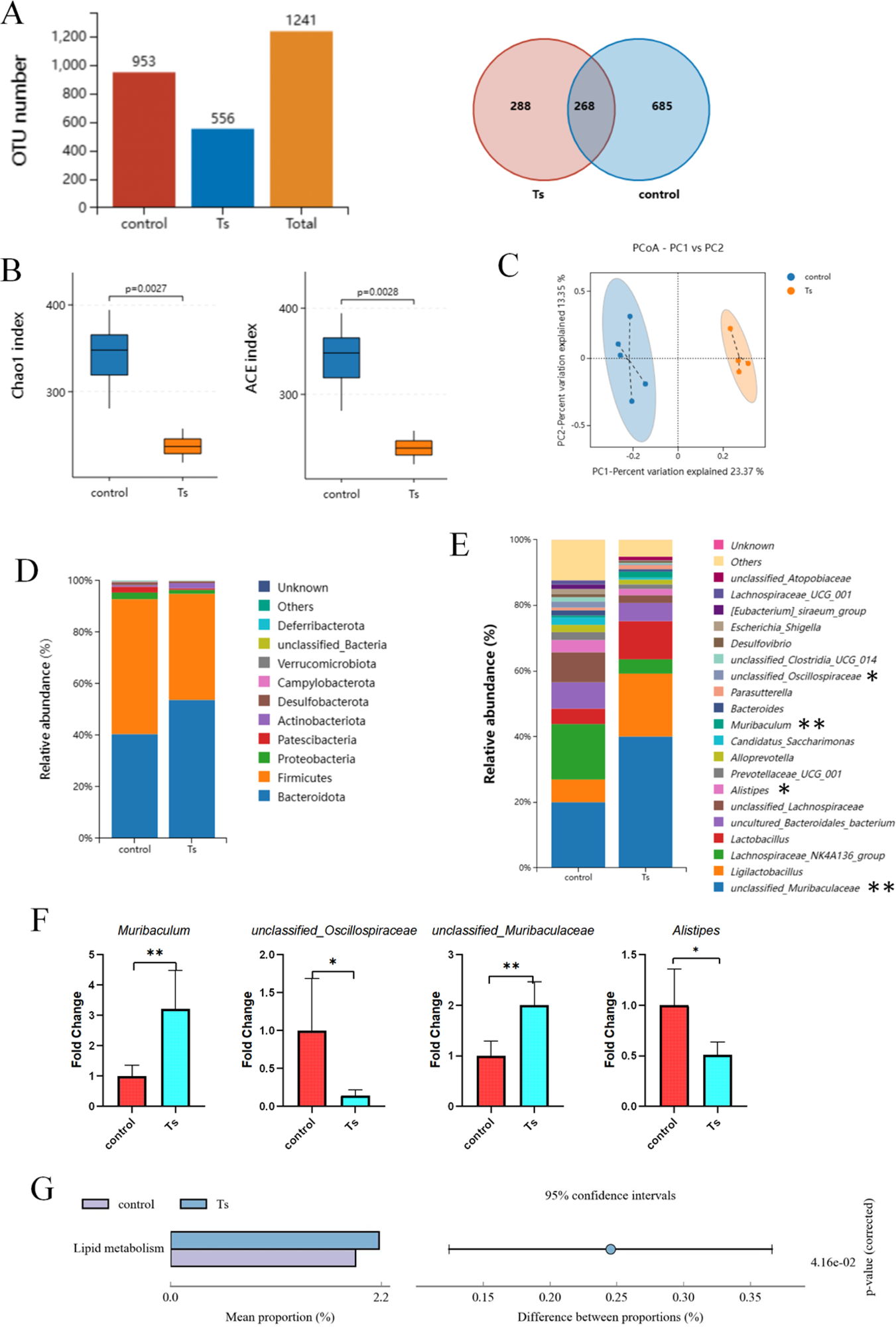
*T. spiralis* infection changes gut microbiota composition in mice. Colon contents were collected and analyzed by using 16S rRNA gene sequencing. (**A**) The number of OTU, (**B**) Alpha diversity analysis (Chao 1 and ACE index), data are shown as Median, maximum, minimum, upper quartile and lower quartile. (**C**) Beta diversity analysis (PCoA based on Binary-Jaccard), (**D**) the relative abundance of OTUs at phylum and (**E**) the relative abundance of the top 20 genus in *T. spiralis*- infected mice and control mice. (**F**) The ratio of *Muribaculum*, *unclassified_Muribaculaceae*, *unclassified_Oscillospiraceae* and *Alistipesin* between groups, data are shown as the means ± SD. \**P* <0.05, \*\**P*<0.001 versus the control group. (**G**) Microbial community functions was predicted by PICRUSt2 using STAMP. Ts: *T. spiralis* infected

### Cohousing with *T. spiralis*- infected mice inhibits DSS-induced colitis

To investigate whether *T. spiralis*- induced alterations in gut microbiota can improve colitis, a cohousing experiment was conducted. Mice were infected with *T. spiralis* for 35 days and then housed together with control mice. DSS-induced colitis was performed after mice were cohoused 4 weeks (**Fig 3A**). Results demonstrated that cohousing had no discernible impact on the development of DSS-induced colitis in mice present *T. spiralis* infection (**S2 Fig**). However, cohousing with *T. spiralis*- infected mice ameliorated the severity of DSS-induced colitis in mice without *T. spiralis*- infected, as evidenced by assessments of body weight, disease activity index, microscopic damage, and histopathology scores (**Fig 3B–3E**).

**Fig 3.**
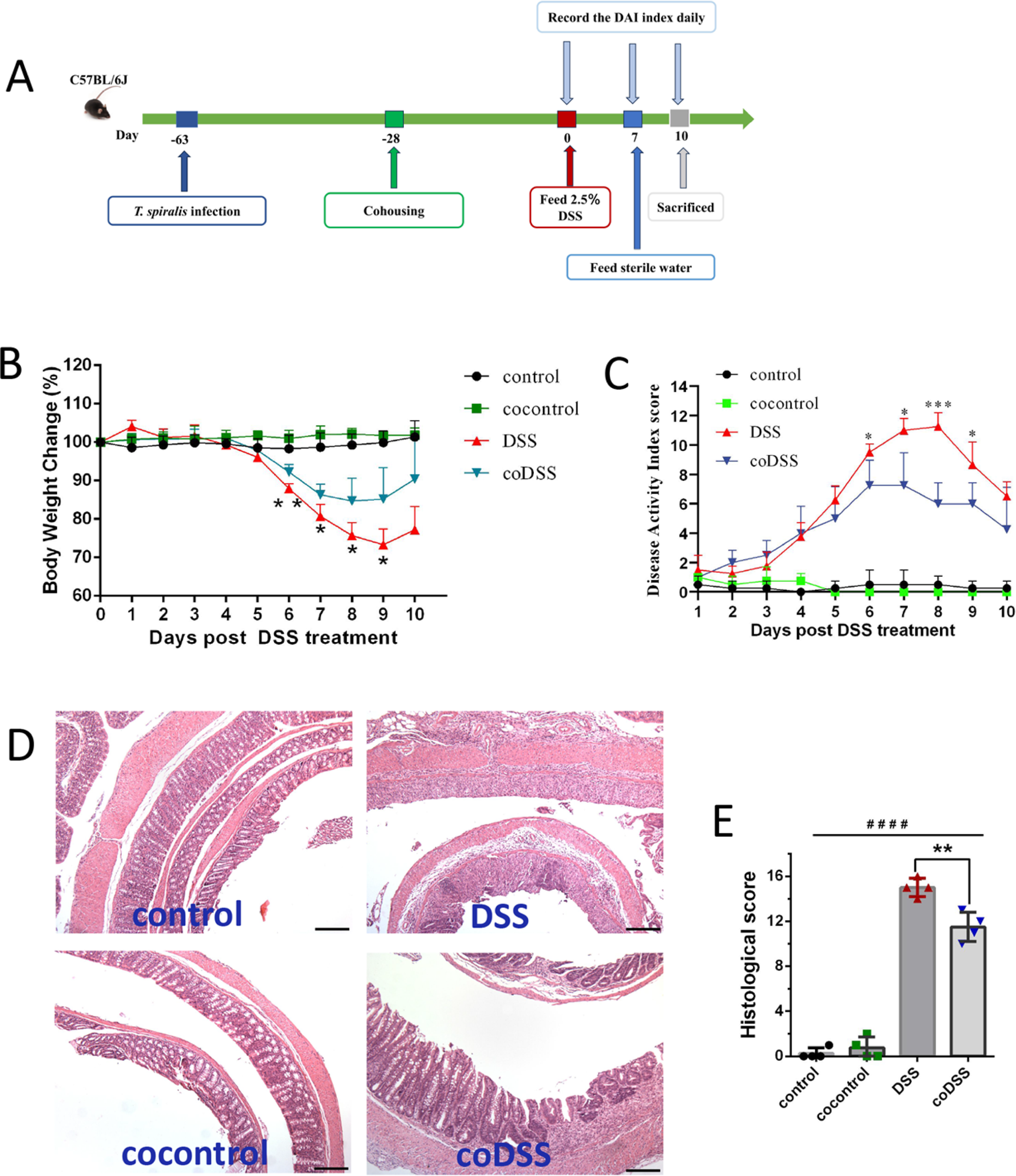
Cohousing with *T. spiralis*- infected mice ameliorates the severity of colitis. **(A)** Infection, cohousing and DSS-induced colitis schedule. The mice were gavaged with *T. spiralis* 5 weeks prior to cohousing, and subsequently induced colitis after 4 weeks of cohousing. (**B**) Weight change in percent. (**C**) The changes in disease activity index (DAI), scored from diarrhea, bleeding and body weight loss. (**D**) The histopathological changes in the colon tissues were examined by H&E staining, the black bar indicates 200 μm. (**E**) Histopathological scores were determined for the colon tissue samples. The data shown are means ± SD. Representative results from one out of two independent experiments with n = 4. \**P* <0.05, \*\**P* <0.01, \*\*\**P* <0.001 versus the DSS group. cocontrol: cohousing with *T. spiralis*- infected mice; DSS: DSS-induced colitis; co-DSS: cohousing with *T. spiralis*- infected mice and DSS-induced colitis

### Cohousing with *T. spiralis*- infected mice alters gut microbiome composition

Subsequently, to investigate the potential impact of cohousing with *T. spiralis*- infected mice on gut microbiota composition, the control mice were housed in the same cage as *T. spiralis* -infected mice for a duration of four weeks. Then, mouse gut microbiota was determined using bacterial 16S rRNA gene sequencing. Results showed that although there was no statistically difference in Chao1 index and ACE index (**Fig 4A**); PCoA shown significantly different clustering of microbiota composition between control mice and cohousing control mice (**Fig 4B**). The data also indicated that mice cohousing with *T. spiralis*- infected mice exhibited similar alterations in the abundance of the bacterial phylum Bacteroidota and Firmicutes and genus *Muribaculum*, *unclassified_Muribaculaceae, unclassified_Oscillospiraceae* and *Alistipes* as those alteration in *T. spiralis*- infected mice (**Fig 4C-4E**). Additionally, there was a decrease in the relative abundances of *unclassified_Clostridia_UCG_014* in cohousing control mice compared to control mice (**Fig 4E**). Furthermore, our data revealed no statistically significant differences in the Chao1 index, ACE index, PCoA, phyla, genera, or cladogram obtained from the LEfSe analysis between cohoused and *T. spiralis*- infected mice (**S3 Fig**). These results, therefore, demonstrate that cohousing with *T. spiralis*- infected mice alters the gut microbiota composition.

**Fig 4.**
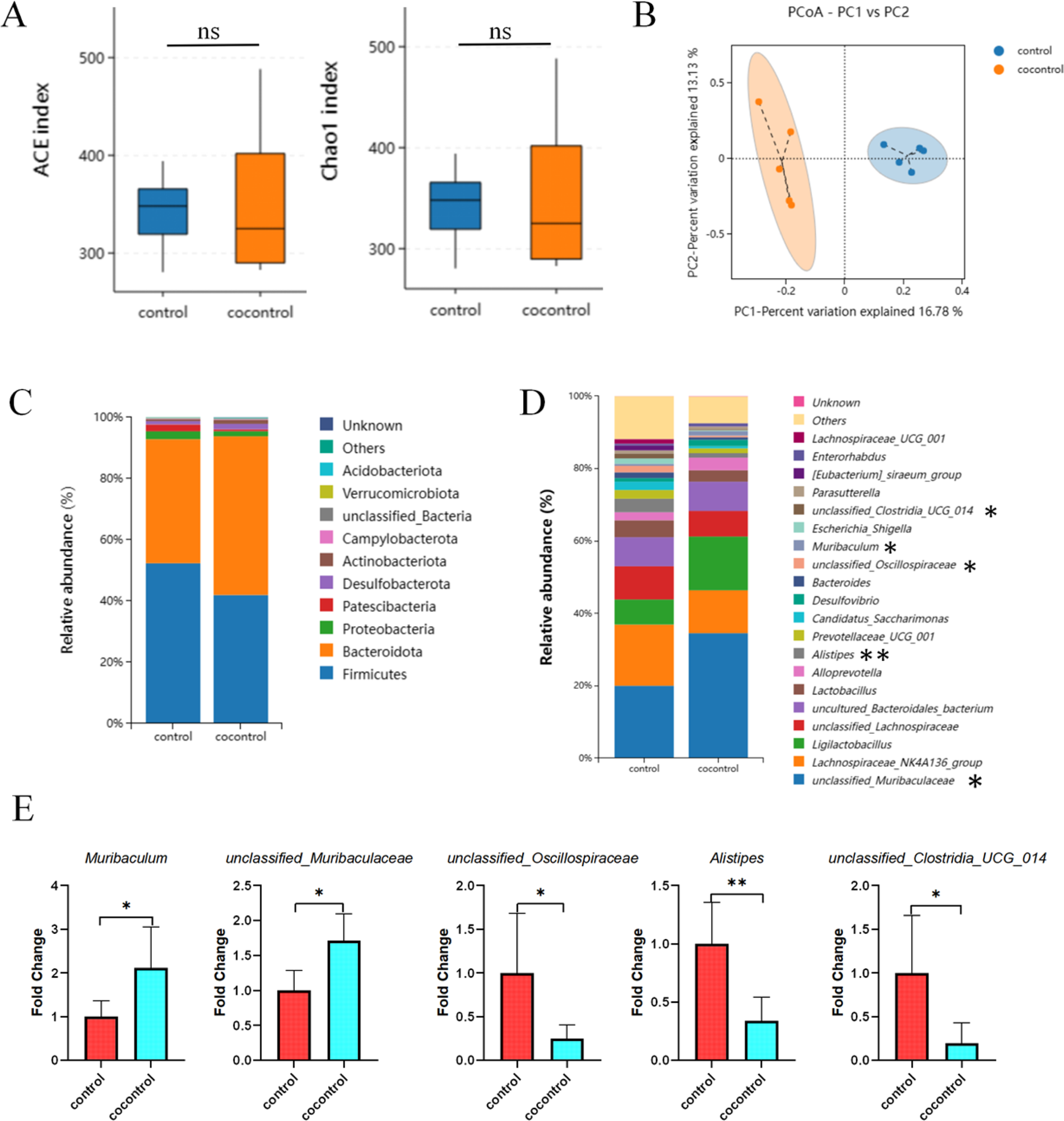
Cohousing with *T. spiralis*- infected mice induces alterations in the composition of the gut microbiome. (**A**) Alpha diversity analysis (Chao 1 and ACE index), data are shown as Median, maximum, minimum, upper quartile and lower quartile. (**B**) Beta diversity analysis (PCoA based on Binary-Jaccard). The relative abundance of the top 10 phyla (**C**) and the top 20 genus (**D**) in mice. (**E**) The ratio of *Muribaculum*, *unclassified_Muribaculaceae*, *unclassified_Oscillospiraceae, Alistipesin* and *unclassified_Clostridia_UCG_014* between groups control mice and mice cohousing with *T. spiralis*- infected mice, data are shown as the means ± SD. \**P* <0.05, \*\**P*<0.001 versus the control group. cocontrol: cohousing with *T. spiralis*- infected mice

### Feces from *T. spiralis*- infected mice increase SCFAs

The alterations in gut microbiota composition are invariably accompanied by changes in metabolites. To explore changes of intestinal metabolome after *T. spiralis* infection, GC-MS was used to analyze short-chain fatty acids (SCFAs) in fecal samples. The PCA score plot and OPLS-DA score plot showed that fecal samples from control mice and *T. spiralis*- infected mice could be easily divided into two distinct clusters (**Fig 5A and 5B**). Based on VIP > 1 and *P* < 0.05 by OPLS-DA model, propionic acid was found as a potential biomarker (**Fig 5C**). Moreover, Permutation Test was performed to verify the validity of OPLS-DA model. And we found that Q2 actually observed was shown to the right of the random distribution and *P* < 0.05, which indicated that the OPLS-DA model had good differential stability and no fitting phenomenon (**Fig 5D**). Furthermore, Support Vector Machines were further employed to validate the difference of SCFAs concentration between control mice and *T. spiralis*- infected mice (**Fig 5E**), revealing an increase in hexanoic acid, isovaleric acid, valeric acid, butyric acid and propionic acid in *T. spiralis*- infected mice compared to control mice (**Fig 5F**). Notably, the difference in propionic acid was found to be statistically significant, with a more than twofold increase observed in *T. spiralis*- infected mice compared to control mice (**Fig 5G and 5H**). Additionally, we also observed that hexanoic acid, isovaleric acid, valeric acid, butyric acid, propionic acid and acetic showed an increase in *T. spiralis*- infected DSS-treated mice compared to DSS-treated mice alone (**Fig 5I**).

**Fig 5.**
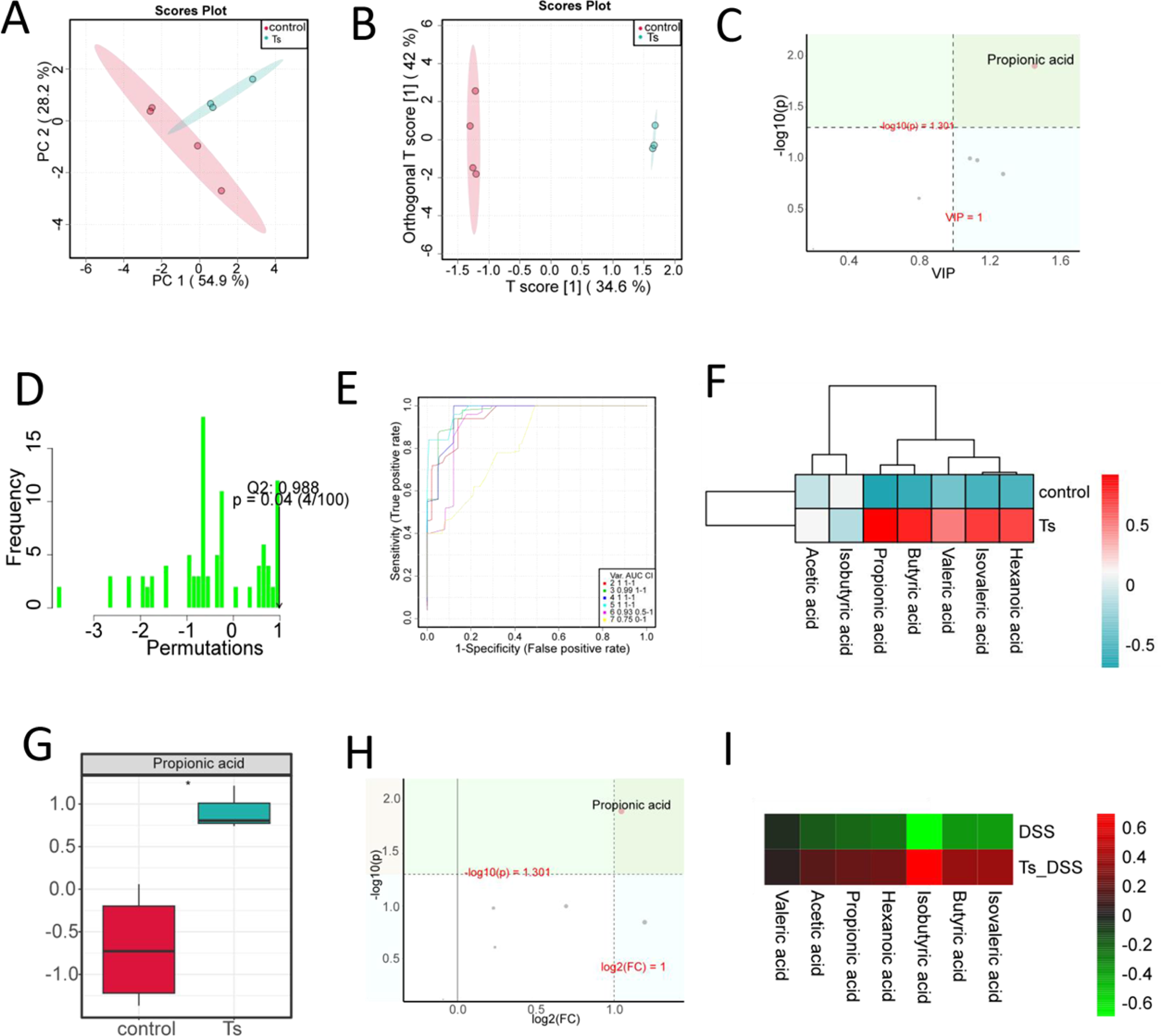
Feces from *T. spiralis*- infected mice contain increase amounts of short-chain fatty acids (SCFAs). (**A**) PCA score plot and (**B**) OPLS-DA score plot shown the difference of control mice and *T. spiralis* -infected mice. (**C**) Volcano map of SCFAs by OPLS-DA model. The abscissa represents Value Importance in Projection (VIP); the ordinate represents the level of significant difference (−Log10 (p)). (**D**) Permutation Test was performed to verify the validity of OPLS-DA model. (**E**) Receiver Operating Characteristic (ROC) curve by Support Vector Machines (SVM). (**F**) Heat-map of SCFAs between control mice and *T. spiralis*- infected mice. (**G**) Box plot of propionic acid, data are shown as Median, maximum, minimum, upper quartile and lower quartile. (**H**) Volcano map of SCFAs. The abscissa represents the logarithm of the relative content fold change (Log2(FC)) of a metabolite in two groups; the ordinate represents the level of significant difference (−Log10(p)). (I) Heat-map of SCFAs between DSS-treated alone group (DSS) mice and *T. spiralis*- infected DSS-treated group (Ts_DSS). \**P* <0.05 compared to control

### *T. spiralis* infection alters immune responses and epithelial barrier properties in DSS-induced colitis mice by modulating the gut microbiome

To explore for the anti-inflammatory role of *T. spiralis* infection or cohousing with infected mice, we investigated tissue immuno-regulatory environment in mice. Lymphocyte were prepared from MLN and stimulated by anti-CD3 antibody. IL-10 in the supernatants were analyzed by ELISA kits. Results showed that the IL-10 production in MLN was upregulated when the mice were induced colitis by DSS in the presence or absence of *T. spiralis* infection, antibiotic treatment, or cohousing (*P* <0.05) (**Fig 6A**). Besides, during DSS-induced colitis, *T. spiralis*- infected mice and cohousing mice exhibited a significantly elevated level of IL-10 production in the MLN compared to DSS-treated mice alone; however, the increase of IL-10 production in *T. spiralis*- infected mice was abolished upon antibiotic treatment (**Fig 6A**). Additionally, the expression level of IL-10, IL-1β and IL-6 were assessed in colon tissue using RT-qPCR. Results revealed an upregulation of IL-10 expression and a downregulation of IL-1β and IL-6 expression in *T. spiralis*- infected mice compared to DSS-treated mice alone during the induction of colitis (**Fig 6B-6D**). However, the differences in the expressions of these cytokines disappeared when the mice were treated with antibiotics or cohoused with *T. spiralis*- infected mice (**Fig 6B-6D**).

**Fig 6.**
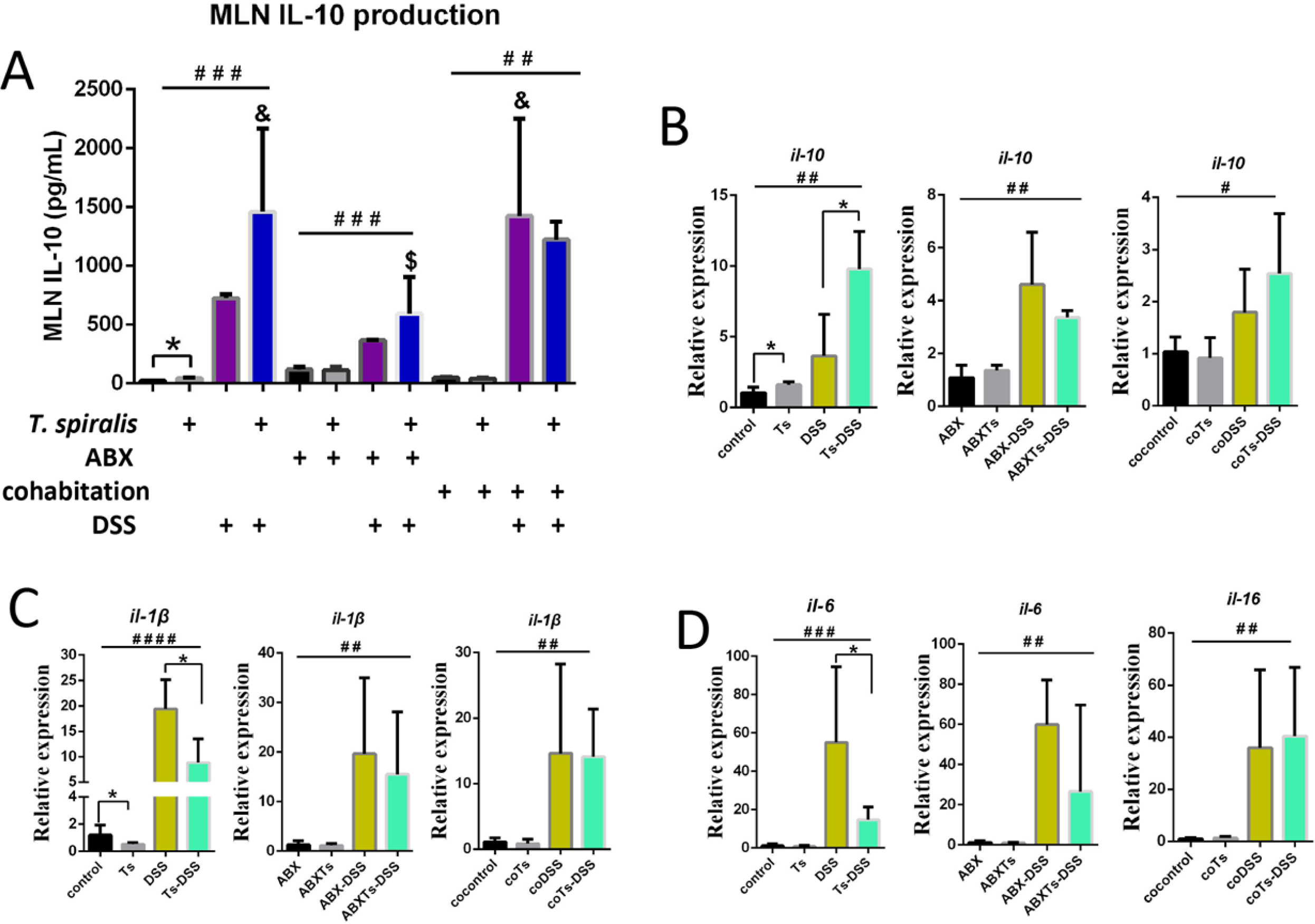
*T. spiralis* infection results in the regulation of both pro-inflammatory and immunoregulatory cytokine responses during DSS-induced colitis by modulating the gut microbiome. Mice infected with *T. spiralis* were subjected to antibiotic treatment or cohousing with control mice, followed by administration with or without DSS. (**A**) Lymphocytes were prepared from MLN and stimulated by anti-CD3 antibody. IL-10 in the supernatants were analyzed by ELISA kits. (**B-D**) The colonic tissues were collected; and the mRNA expression of IL-10, IL-1β and IL-6 was determined using RT-qPCR. The data shown are means ± SD (n = 4-5 mice per group) from one of three experiments performed showing similar results. \**P* <0.05 between the indicated groups; #*P* <0.05, ##*P* <0.01, ###*P* <0.001, ####*P* <0.0001 versus the respective control group; & *P* <0.05 versus the DSS group; $ *P* <0.05 versus the Ts-DSS group. ABX: antibiotic treated; Ts, *T. spiralis* infected; ABXTs: *T. spiralis* infected and antibiotic treated; DSS: DSS induced colitis; Ts-DSS: *T. spiralis* infected and DSS induced colitis; ABX-DSS: antibiotic treated and DSS induced colitis; ABXTs-DSS: *T. spiralis* infected, antibiotic treated and DSS induced colitis; cocontrol: cohousing with *T. spiralis* infected mice; coTs: *T. spiralis* infected and cohoused; coDSS: cohousing with *T. spiralis* infected mice and DSS induced colitis; coTs-DSS: *T. spiralis* infected, cohoused and DSS induced colitis

To further determine whether *T. spiralis* infection could affect colonic goblet cell response, contributing to the observed protection, Periodic Acid Schiff (PAS) staining was performed on the mouse colonic sections. Our results demonstrated a significant decrease in the number of goblet cells (PAS+ cells) in the colons of DSS-treated mice compared to their respectively control mice (*P* <0.05) (**Fig 7**). During DSS-induced colitis, *T. spiralis*- infection and cohousing resulted in a significant increase in goblet cell numbers compared to DSS-treated mice alone; however, this increase was abolished upon antibiotic treatment (**Fig 7A and 7B**). Furthermore, administration of antibiotics resulted in a decrease in the number of colonic goblet cells in mice infected with *T. spiralis* compared to those receiving antibiotic treatment alone. Taken together, these results demonstrate that *T. spiralis* infection alters immune responses and epithelial barrier properties in DSS-induced colitis mice by modulating the gut microbiome.

**Fig 7.**
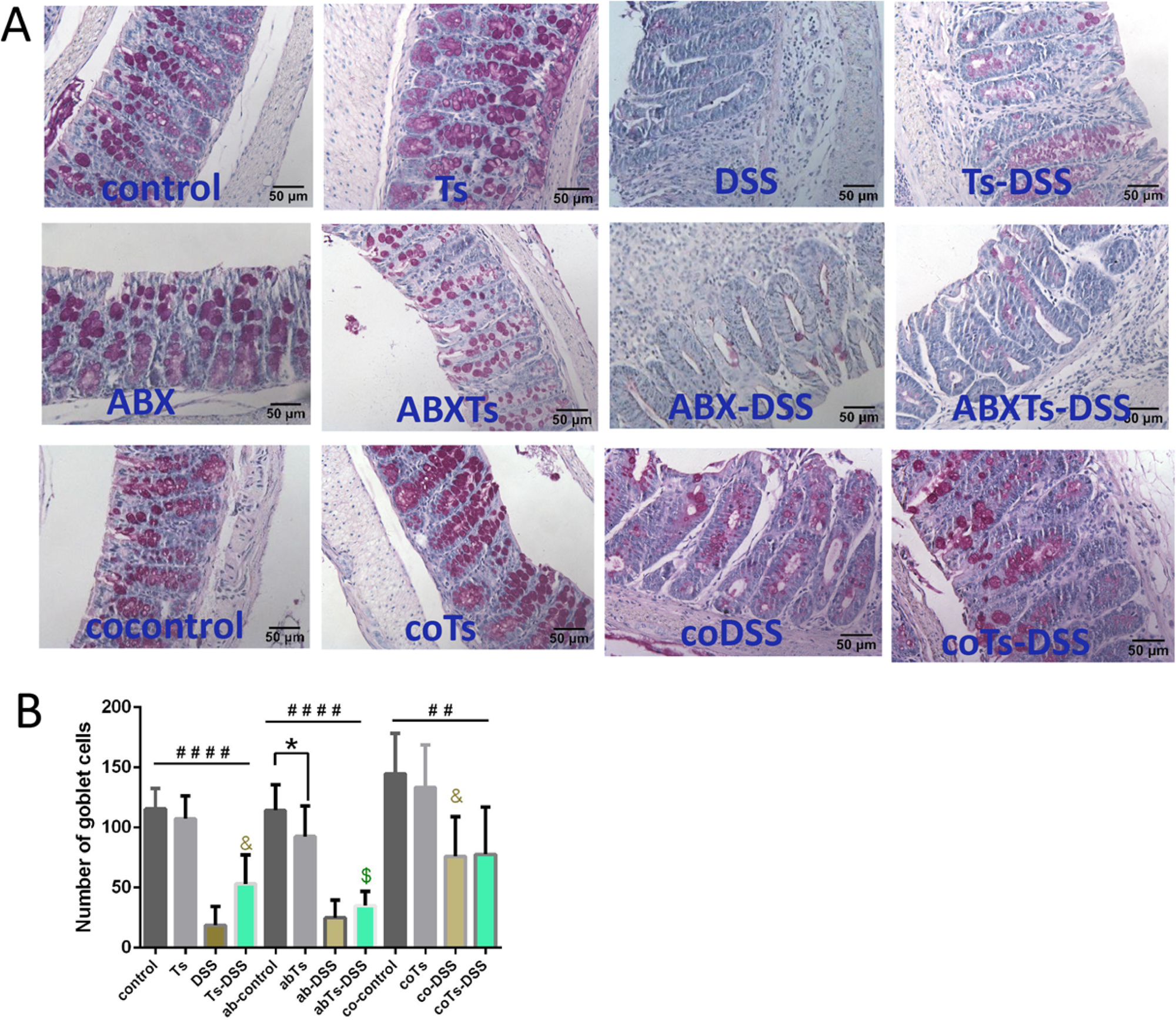
*T. spira*lis infection modulates epithelial barrier properties during DSS-induced colitis by modulating the gut microbiome. Mice infected with *T. spiralis* were subjected to antibiotic treatment or cohousing. (**A**) Goblet cells were stained with periodic acid-schiff (PAS), and (**B**) the number of goblet cells was calculated as the average score of 10 random fields from each mouse. The data shown are means ± SD (n = 4-5 mice per group). \**P* < 0.05 between the indicated groups; ##*P* < 0.01, ####*P* < 0.0001 versus the respective control group; & *P* < 0.05 versus the DSS group; $ *P* < 0.05 versus the Ts-DSS group. ABX: antibiotic treated; Ts, *T. spiralis* infected; ABXTs: *T. spiralis* infected and antibiotic treated; DSS: DSS induced colitis; Ts-DSS: *T. spiralis* infected and DSS induced colitis; ABX-DSS: antibiotic treated and DSS induced colitis; ABXTs-DSS: *T. spiralis* infected, antibiotic treated and DSS induced colitis; cocontrol: cohousing with *T. spiralis* infected mice; coTs: *T. spiralis* infected and cohoused; coDSS: cohousing with *T. spiralis* infected mice and DSS induced colitis; coTs-DSS: *T. spiralis* infected, cohoused and DSS induced colitis

## Discussion

Helminth infections are known to be powerful modulators of the human immune response, and numerous studies now highlight the effects may have on human infectious, inflammatory, and metabolic diseases [14]. The helminth *Trichinella spiralis* is the causative agent of trichinosis. Humans contract *T. spiralis* by consuming uncooked meat that carries encysted *T. spiralis* larvae in muscle tissue; then larvae are released by gastric fluids, molt and mature into adult worm. The female adult worms release newborn larvae after copulation, which travel through the circulatory system to reach skeletal muscle cells [14, 15]. As a helminth with all stages of larval and adult development occurring within the same host organism, *T. spiralis* is one of the most successful parasitic symbiotes, ensuring its survival and immune dialogue with the host [16]. Multiple studies have confirmed the anti-inflammatory effect of *T. spiralis* in animal models. Chronic infection with *T. spiralis* can mediate protection against allergic asthma, *Pseudomonas aeruginosa*-induced pneumonia, influenza or respiratory syncytial virus-associated pathologies in mice [15, 17–19]. Evidence also indicates that *T. spiralis* infection attenuates *Citrobacter rodentium*, acetic acid, trinitrobenzesulfonic acid or DSS-induced colonic damage [8, 20–23]. However, the potential influence and mechanism of *T. spiralis* infection inducing alterations in the gut microbiota on the development of IBD is still largely undetermined. In this study, we analyzed the intersection of *T. spiralis* and gut bacteria in the regulation of colitis and related mechanisms by infecting mice with *T. spiralis*, administering broad-spectrum antibiotics to mice, or cohousing mice with *T. spiralis*-infected counterparts. Our findings revealed that prior infection with *T. spiralis* or cohousing with *T. spiralis*- infected mice could ameliorate DSS-induced colitis through an increase abundance of SCFA-producing bacteria. Consistent with previous effects, our current study demonstrated that infection with *T. spiralis* significantly reduced disease activity index, ameliorated clinical symptoms, and improved colonic histological damage in mice with DSS-induced colitis. However, treatment with broad spectrum antibiotics was found to prevent the inhibition of colitis evoked by *T. spiralis* infection, suggesting that lack of inhibition of colitis was linked to the microbiota. Subsequently, we employed 16S rRNA gene sequencing to investigate the impact of *T. spiralis* infection on gut microbiota and observed significant alterations in both diversity and composition. *T. spiralis* infection led to an increase in the abundance of Bacteroidota while decreasing the abundance of Firmicutes at the phylum level. Specifically, characterized by an increase in the abundance of the bacterial genus *Muribaculum* and *unclassified_Muribaculaceae and* a decrease in the abundance of the opportunistic pathogens [24, 25] bacterial genus *unclassified_Oscillospiraceae* and *Alistipes* in the top 20 genera. It is worth noting that the differential effects of feces from *T. spiralis*- infected mice may also arise from variations in gut bacteria composition due to factors such as animal source, housing conditions, diet or duration of infection [26–28]. Dissecting the role of the microbiota in the anti-colitic evoked by *T. spiralis* infection, cohousing the control mice with *T. spiralis* infection mice conferred protection from DSS-induced colitis. Furthermore, cohousing control mice exhibited similar changes in the bacterial genera *Muribaculum, unclassified_Muribulaceae, unclassified_Oscillospiraceae*, and *Alistipes* as those observed in *T. spiralis*- infected mice. *Muribaculum* is a major forager of mucin monosaccharides, which could impede the colonization of *Clostridioides difficile* [29]. The infection of *T. spiralis* significantly enhanced the relative abundance of *Muribaculum*, thereby potentially impeding pathogen colonization and maintaining intestinal homeostasis simultaneously. Meanwhile, both *Muribaculum* and *unclassified_Muribaculaceae* belong to the family Muribaculaceae [30], whose abundance exhibits a negative correlation with gut inflammation [31–33]. Our observations align with previous studies showing a robust interaction between *T. spiralis*- modulated *Muribaculum* and *unclassified_Muribulaceae* and IBD. In addition to taxonomic composition, the functional profiling of microbial communities revealed a significant enhancement in lipid metabolism function within gut bacteria derived from *T. spiralis*- infected mice.

SCFAs are metabolized by intestinal bacteria from a fiber-rich diet that is otherwise indigestible, and primarily consist of acetic acid, propionic acid and butyric acid [34]. A growing body of research has indicated that SCFAs play critical roles in IBD [35–38]. Our results provide evidence indicating that *T. spiralis*- enriched *Muribaculum* and *unclassified_Muribulaceae* are correlated with enhanced production of SCFAs [30], especially propionic acid, which has been demonstrated to possess functions in reducing the relapse rate and disability progression of multiple sclerosis by increasing expression of Treg-cell-inducing genes, such as IL-10 [39], as well as alleviating intestinal inflammation by enhancing goblet cell differentiation and mucus function [40].

IL-10 is a pre-dominantly anti-inflammatory cytokine with an essential role in maintaining gastrointestinal homeostasis, which potently inhibits production of most inducible chemokines that are involved in inflammation [41]. Several helminthes attenuate colitis in DSS models by inducing anti-inflammatory cytokine expression and downregulating pro-inflammatory cytokines expression [7, 9]. Consistent with previous studies, helminth *T. spiralis* demonstrated a significant upregulation of IL-10 and downregulation of IL-1β and IL-6. Interestingly, mice cohousing with *T. spiralis*- infected mice also exhibited a significantly heightened level of IL-10 production, while the discrepancy in IL-10 production was eliminated upon administration of antibiotics to the mice. These findings suggest a robust interplay between *T. spiralis*- modulated microbiota and anti-inflammatory cytokine IL-10.

Goblet cells are recognized as key regulators of intestinal barrier homeostasis and play a crucial role in defending against enteric pathogen infections [42]. Alternations of goblet cell number and mucus formation are associated with the pathogenesis and progression of IBD, including ulcerative colitis and Crohn’s disease [40]. In this study, we identified the critical role of *T. spiralis*- modulated microbiota in alleviating DSS-induced colitis and enhancing goblet cell proportions. During DSS-induced colitis, *T. spiralis* infection and cohousing resulted in a significant increase in goblet cells; however, this increase was abolished after antibiotic treatment. These findings suggest a robust interplay between *T. spiralis*- modulated microbiota and goblet cell proportions. However, the role of SCFAs in mediating the beneficial effects of *T. spiralis*- modulated microbiota on colitis remains a crucial question that requires further investigation.

In conclusion, we revealed the protective effect of a prior infection with *T. spiralis* on DSS- induced colitis in mice. Specifically, *T. spiralis* infection reshaped the gut microbiome of mice, resulting in an increased abundance of SCFA-producing bacteria *Muribaculum* and *unclassified_Muribaculaceae* and thereby producing a larger amount of propionic acid. Furthermore, the gut microbiome modulated by *T. spiralis* exhibited enhanced goblet cell differentiation and elevated IL-10 levels, ultimately ameliorating experimental colitis. These findings suggest that the modulation of host microbiota during *T. spiralis* infection plays a crucial role in the suppression of colitis. Our research gives beneficial insights into the processes by which helminth alleviates colitis.

## Materials and methods

### Experimental animals

Pathogen-free, 6-week-old female C57BL/6J mice were purchased from GemPharmatech (Jiangsu, China). They were fed autoclaved food and water. All of the animal experiments were approved by the Zhengzhou University Life Science Ethics Committee (No. ZZUIRB GZR 2021–0044).

### Helminth infection and DSS-Induced Colitis

The helminth parasite *T. spiralis* (ISS534) was collected as previously described [43]. Four groups of mice (5 mice per group, include control: PBS control group, Ts: *T. spiralis-* infected group, DSS: DSS-induced colitis group, and Ts-DSS: *T. spiralis-* infected DSS-induced colitis group) were used. Three independent experiments were performed. *T. spiralis* infected groups were administered 100 muscle larvae orally by gavage with a 21-gauge feeding needle. Control group was gavaged with PBS at the same time. *T. spiralis* infection was incubated for a period of 35 days prior to treatment of the mice with DSS. Experimental colitis was induced as previous study[43]. A 2.5% (wt/vol) solution of DSS (M.W. 36000–50000 kDa; MP Biomedicals, Cat. no. 106110) were administered to mice as a substitute for autoclaved water for 7 days. At day 7 post DSS administration, the DSS solution was replaced by autoclaved water. After a further 3 days of water fed, mice were sacrificed and samples were obtained and processed. Clinical parameters including body weight, bleeding, and stool consistency were recorded daily to calculate the clinical disease score (disease activity index, DAI) [44].

### Antibiotic treatment of mice

Mice were infected with *T. spiralis* for 35 days, then treated with a broad-spectrum cocktail of antibiotics (ABX: drinking water, ampicillin (1g/L), vancomycin (0.5g/L), neomycin (1g/L), and metronidazole (1g/L)) daily for 14 days to deplete the gut microbiota. To evaluate the role of gut microbiota in helminth-ameliorated colitis, four groups of mice (3-4 mice per group, include ABX: antibiotic treatment control group, abTs: *T. spiralis-* infected antibiotic-treated group, ab-DSS: antibiotic-treated DSS-induced colitis group, and abTs-DSS: *T. spiralis-* infected antibiotic-treated and DSS-induced colitis group) were used. Two independent experiments were performed. DSS induced colitis was performed 2 weeks after antibiotic treatment.

### Cohousing of mice

To investigate whether helminth-induced alterations in gut microbiota can improve colitis, a cohousing experiment was conducted between helminth-infected mice and control mice. Mice were infected with *T. spiralis* for 35 days and then housed together with PBS-treated control mice. Four groups of mice (4 mice per group, include cocontrol: cohoused control group, coTs: *T. spiralis* infected and co-housed group, co-DSS: cohoused and DSS-induced colitis group, and coTs-DSS: *T. spiralis-*infected, cohoused, and DSS-induced colitis group) were used. Two independent experiments were performed. DSS induced colitis was performed 4 weeks after cohousing.

### Histopathological examinations

A 0.5-1.0 cm colon piece (cecum side) was collected for the measurement of tissue cytokine expression by using RT-qPCR, the rest of the colon tissue was cut longitudinally and rolled into the shape of a Swiss roll. The colons were fixed in 4% formaldehyde and embedded in paraffin. The process tissues were sectioned into 5 μm thick slices and stained with hematoxylin and eosin (H&E). The extent of colonic inflammatory damage was assessed using the scoring system described by Wang et al [45]. Goblet cells were stained with periodic acid-schiff (PAS) following manufacturer instructions, and the number of goblet cells per field in the colon were quantified.

### Lymphocyte isolation and IL-10 production measurement

Mice were sacrificed 10 days after DSS treatment. Lymphocyte suspensions were prepared from mesenteric lymph nodes (MLN) by pressing the cells through a 70-μM nylon cell strainer (Falcon; BD Labware) in complete Dulbecco’s modifed Eagle’s medium (cDMEM) (10% fetal calf serum, 2mM L-glutamine, 100U of penicillin/mL, 100μg of streptomycin/mL, 1% nonessential amino acids, and 110mg/L sodium pyruvate, 4.5g/L D-Glucose). Red blood cells were lysed. Cells (1x10^6^ cells/mL) were cultured in a 48-well plate pre-coated with anti-CD3 monoclonal antibody (BD Pharmingen, 5μg/mL), and culture supernatants were collected after 72h of stimulation and stored at -80°C until assayed for cytokine production. IL-10 in the supernatants were analyzed by ELISA kits (Proteintech, KE10008).

### RNA extraction and RT-qPCR

RNA extraction was performed by lysing 50 mg of mouse colon tissue samples with Trizol reagent (Invitrogen, USA) and then converted to cDNA. Quantitative real-time reverse-transcription PCR (RT–qPCR) assays were performed on a 7500 Fast Real-Time PCR system (Applied Biosystems, USA). The primer sequences were listed in **S1 Table**. The relative mRNA expression levels of the target genes were normalized to those of the indicated housekeeping gene (GAPDH) and were quantified using the comparative Ct method and the formula 2^-ΔΔCT^.

### Gut microbiome analysis

Fecal samples were collected from the control mice, *T. spiralis* infected mice and cohousing mice. DNA was extracted from fecal samples and added spike-in DNA at a certain ratio. Based on 16s v3+v4_b, primes were designed as F: ACTCCTACGGGAGGCAGCA; R: GGACTACHVGGGTWTCTAAT and the sequencing adapter was added at the end of the primers. PCR amplification was carried out, and the products were purified, quantified and normalized to form a sequencing library. Qualified libraries were sequenced using Illumina Novaseq 6000 for paired-end reads. The raw reads were preconditioned using Trimmomatic v0.33, cutadapt 1.9.1, Usearch v10 and so on to get high-quality reads. Operational taxonomic units (OTUs) based on ≥97% sequence similarity were obtained by using the dada2 method included in the QIIME2 software. Alpha diversity (Chao 1 and ACE index) and Beta diversity (PCoA based on Binary-Jaccard) were analyzed by using QIIME2 software. Using SILVA as a reference database and Naive Bayesian classifier for taxonomic annotation of feature sequences.

### Measurement of fecal short-chain fatty acids (SCFAs)

Fecal samples were thawed on ice. 200 mg of sample was suspended with 50 uL of 20% phosphoric acid, added 4-methylvaleric acid to a final concentration of 500ug/mL as internal standard, mixed for 2 min and centrifuged at 14000 g for 20 min. Su pernatant was analyzed by Gas Chromatography-Mass Spectrometry (GC-MS). Metabolites were separated on gas chromatography system with Agilent DB-FFAP capillary column (30m × 250um × 0.25um) and then performed mass spectrometry by 5977B MSD Agilent. Area of chromatographicpeak and retention time were extracted by MSD ChemStation software. The concentration of SCFAs was calculated by drawing standard curve. This metabolome analysis workflow was based on MetaboAnalystR [46].

### Statistical analysis

All data were statistically analyzed using GraphPad Prism 8 and presented as mean ± SD. Statistical differences were determined using a two-tailed Student t test or One-way ANOVA with SPSS 21.0 software. A *P* value < 0.05 was considered significant.

### Supporting information

S1 Table Primers used for qRT-PCR analysis

## Data Availability

The datasets used and analyzed in this article are available from the corresponding author upon reasonable request.

## Author Contributions

Study conception and design: SRL, ZQW, JC and HS. Acquisition of data: MJ, HRZ, ZXL and JJW. Analysis and interpretation of data: SRL, MJ and HS. Drafting of manuscript: SRL, MJ and HRZ. Critical revision of manuscript: ZQW, JC and HS. All authors have read and agreed to the published version of the manuscript.

---

**S1 Fig.**
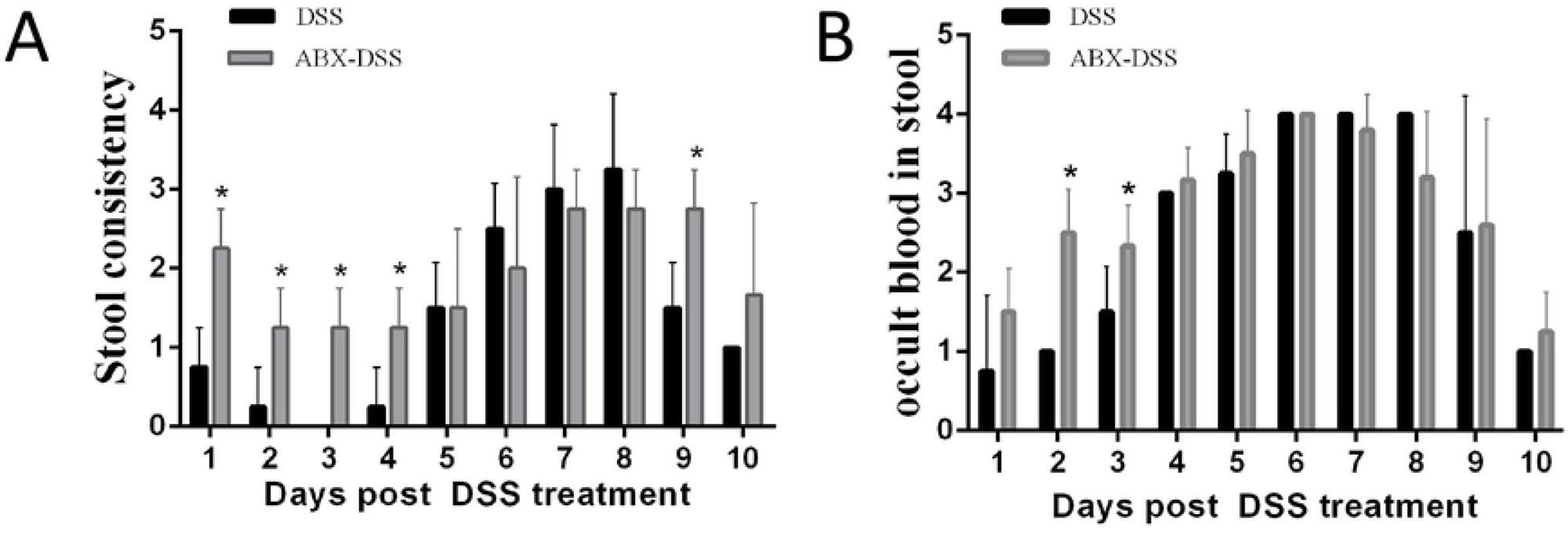
Mice present ABX treatment during the induction of colitis exhibits an earlier onset of (A) diarrhea and (B) rectal bleeding compared to mice without ABX treatment. The data shown are means ± SD. Representative results from one out of two independent experiments with n = 5. \**P*< 0.05 compared to DSS group. DSS: DSS-induced colitis; ABX-DSS: antibiotic-treated and DSS-induced colitis

**S2 Fig.**
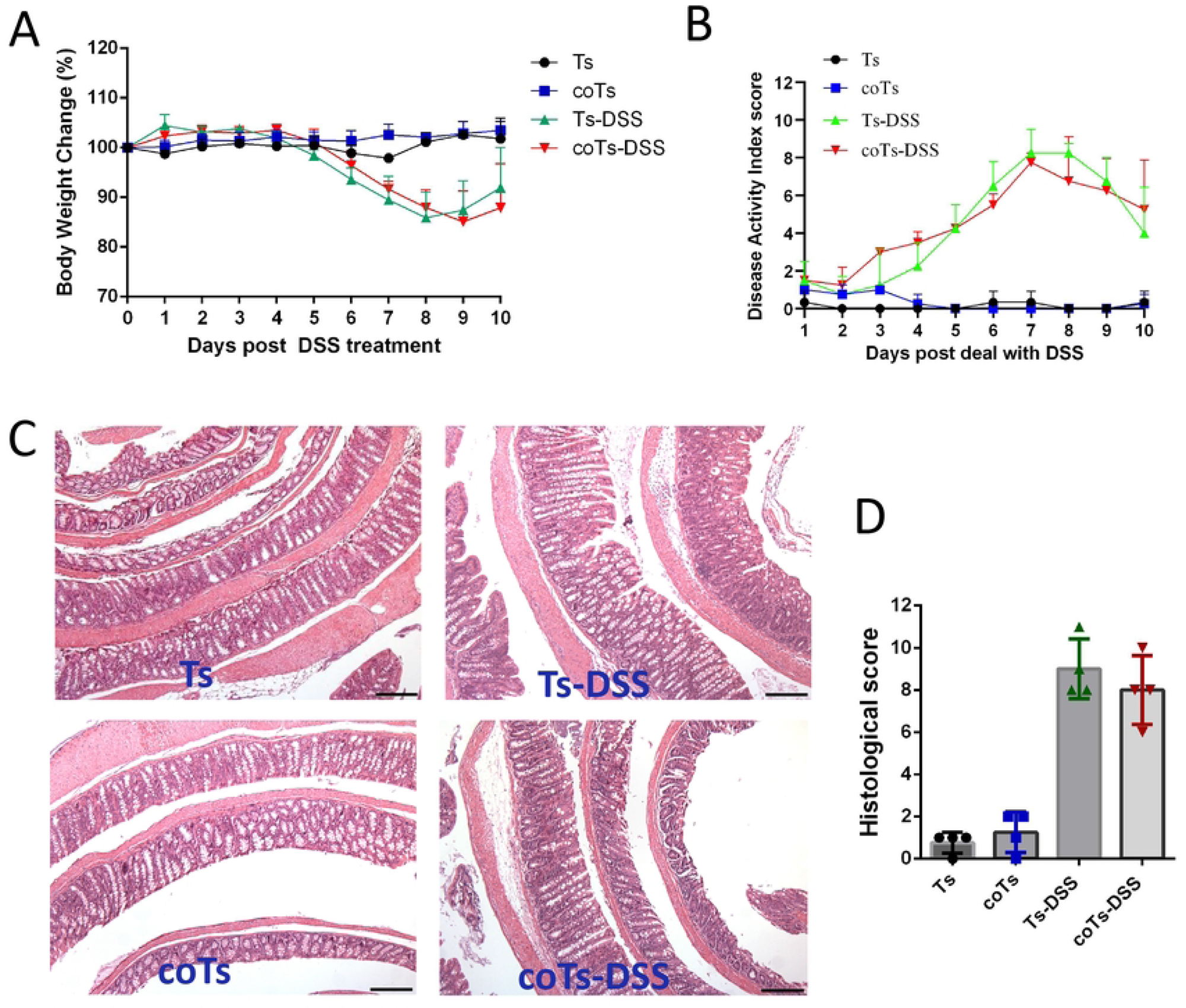
Cohousing has no impact on the DSS-induced colitis in mice present *T. spiralis* infection. (**A**) Weight change in percent. (**B**) The changes in disease activity index (DAI), scored from diarrhea, bleeding and body weight loss. (**C**) The histopathological changes were examined by H&E staining, the black bar indicates 200 μm. (**D**) Histopathological scores. The data shown are means ± SD. Representative results from one out of two independent experiments with n = 4. Ts, *T. spiralis-* infected; coTs: *T. spiralis-* infected and cohoused; Ts-DSS: *T. spiralis-* infected and DSS-induced colitis; coTs-DSS: *T. spiralis-* infected, cohoused and DSS-induced colitis

**S3 Fig.**
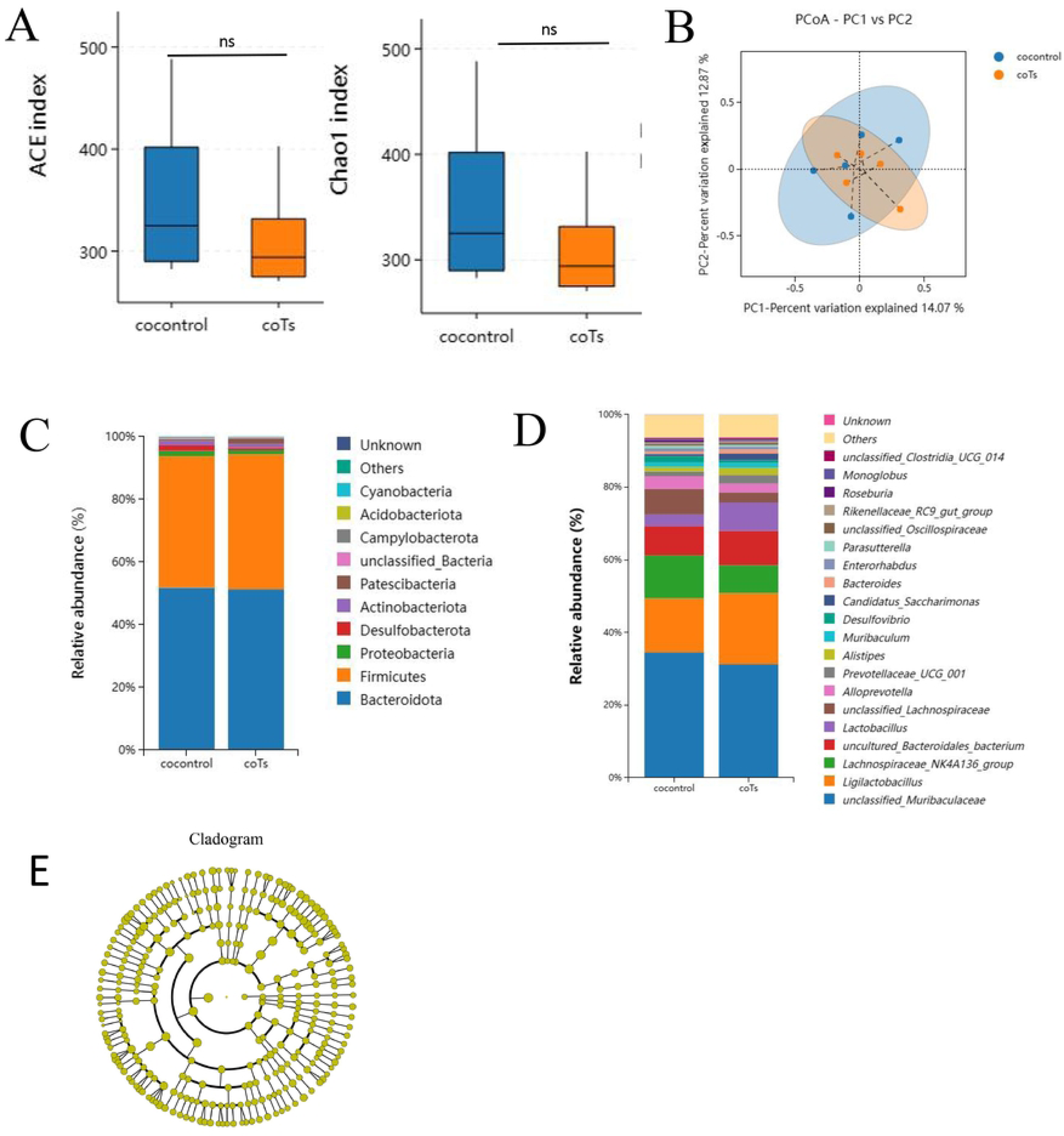
The gut microbiota composition in mice becomes consistent when cohousing of control and *T. spiralis*- infected mice. (**A**) Alpha diversity analysis (Chao 1 and ACE index), data are shown as median, maximum, minimum, upper quartile and lower quartile. (**B**) Beta diversity analysis (PCoA based on Binary-Jaccard). The relative abundance of the top 10 phyla (**C**) and the top 20 genus (**D**) in cohousing control mice and cohousing *T. spiralis*- infected mice. (**E**) Cladogram was obtained from the LEfSe analysis when the effect size threshold of LDA was set to 3.5. cocontrol: cohousing with *T. spiralis*- infected mice; coTs: *T. spiralis* infected and cohoused

